# A level adjusted cochlear frequency-to-place map for estimating tonotopic frequency mismatch with a cochlear implant

**DOI:** 10.1101/2024.06.26.600724

**Authors:** Elad Sagi, Mario A. Svirsky

**Affiliations:** Department of Otolaryngology — Head & Neck Surgery, NYU Grossman School of Medicine, New York, NY

## Abstract

**Objectives:** To provide a level-adjusted correction to the current standard relating anatomical cochlear place to characteristic frequency in humans, and to re-evaluate anatomical frequency mismatch in cochlear implant (CI) recipients considering this correction. It is hypothesized that a level-adjusted place-frequency function may represent a more accurate tonotopic benchmark for CIs in comparison to the current standard.

**Design:** The present analytical study compiled data from fifteen previous animal studies that reported iso-intensity responses from cochlear structures at different stimulation levels. Extracted outcome measures were characteristic frequencies and centroid-based best frequencies at 70 dB SPL input from 47 specimens spanning a broad range of cochlear locations. A simple relationship was used to transform these measures to human estimates of characteristic and best frequencies, and non-linear regression was applied to these estimates to determine how the standard human place-frequency function should be adjusted to reflect best frequency rather than characteristic frequency. The proposed level-adjusted correction was then compared to average place-frequency positions of commonly used CI devices when programmed with clinical settings.

**Results:** The present study showed that the best frequency at 70 dB SPL (BF70) tends to shift away from characteristic frequency (CF). The amount of shift was statistically significant (signed-rank test z = 5.143, p < 0.001), but the amount and direction of shift depended on cochlear location. At cochlear locations up to 600° from the base, BF70 shifted downwards in frequency relative to CF by about 4 semitones on average. Beyond 600° from the base, BF70 shifted upwards in frequency relative to CF by about 6 semitones on average. In terms of spread (90% prediction interval), the amount of shift between CF and BF70 varied from relatively no shift to nearly an octave of shift. With the new level-adjusted frequency-place function, the amount of anatomical frequency mismatch for devices programmed with standard of care settings is less extreme than originally thought, and may be nonexistent for all but the most apical electrodes.

**Conclusions:** The present study validates the current standard for relating cochlear place to characteristic frequency, and introduces a level-adjusted correction for how best frequency shifts away from characteristic frequency at moderately loud stimulation levels. This correction may represent a more accurate tonotopic reference for CIs. To the extent that it does, its implementation may potentially enhance perceptual accommodation and speech understanding in CI users, thereby improving CI outcomes and contributing to advancements in the programming and clinical management of CIs.

## INTRODUCTION

Cochlear implants (CIs) are auditory prosthetic devices that restore hearing to individuals with severe-to-profound hearing impairment. Although a life changing intervention, CIs provide listeners with a distorted input that requires time to perceptually accommodate. For example, CI users may require months to years before achieving asymptotic levels of speech understanding with their clinically programmed settings (Holden et al., 2013; Cusumano et al., 2017; James et al., 2019; Caswell-Midwinter et al., 2022). At least for postlingually deafened individuals, one of many limitations thought to contribute to these distortions is an anatomical place-frequency mismatch. CIs are not typically inserted fully into the cochlea, yet are programmed to provide a range of frequencies thought to be most important for speech understanding (e.g. 200Hz to 8 kHz). The result is a mismatch between frequencies delivered to electrodes, and the tonotopic frequencies associated with the neural elements stimulated by those electrodes prior to losing hearing. By one estimate, this anatomical frequency mismatch can be an octave or more (Landsberger et al., 2015). Although CI users can perceptually adapt to this mismatch over time, this adaptation may be incomplete leaving a perceptual frequency mismatch, which can hinder speech understanding (Canfarotta et al., 2022; Mertens et al., 2022; Dillon et al., 2023; Weller et al., 2023; Dessard et al., 2024; but see Sturm et al., 2024). Alternatively, it has been suggested that programming CIs with a tonotopically correct frequency-to-electrode allocation may provide more benefit than standard-of-care settings, whether in terms of initial fitting to provide a better basis for adaptation (Stakhovskaya et al., 2007; Svirsky et al., 2015), or in terms of better pitch perception (Oxenham and Bernstein, 2004; Jiam et al., 2019) and music perception (Limb and Roy, 2014), or in terms of better electroacoustic integration in EAS CI users (Fu et al., 2017; Dillon et al., 2022; Dillon et al., 2023) and better binaural benefit in bimodal (Pieper et al., 2022), SSD CI (Wess et al., 2017; Sagi et al., 2021), and in bilateral CI users (Xu et al., 2020; Pieper et al., 2022; Kurtz et al., 2023).

At the core of evaluating frequency mismatch in CIs (and their tonotopic correction) is the need for an accurate representation of the cochlear place-frequency map. The most commonly used estimate for the human tonotopic map is that of Greenwood (1961, 1990), originally derived from a consideration of perceptual critical bands and further validated with physiological measures obtained from several species, including humans. Greenwood’s equation takes the following forms:

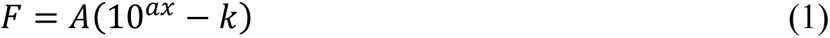

or,

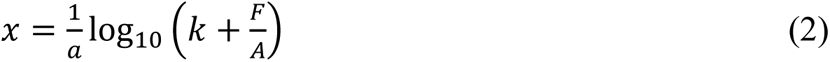

where x is distance along the cochlear partition from the apex, F is frequency (in Hz), and where 𝑎, 𝑘, and 𝐴 are constants. When x is expressed as a proportion of cochlear length (ranging from 0 at the apex to 1 at the base), a common value of 𝑎 = 2.1 was found to provide a consistent fit to frequency-place measures across several species including humans, suggesting that longitudinal cochlear position relative to cochlear length is scale-related across species. The constant 𝑘 varied slightly across species with values between 0.8 and 1. In contrast, the constant 𝐴 varied widely across species, e.g. 𝐴 = 165.4 in humans (Greenwood, 1990) and 𝐴 = 456 in cats (Liberman, 1982), and relates to the upper frequency limit assigned to the most basal location. Commonly used coefficient values for humans in the CI literature are 𝐴 = 165.4, 𝑎 = 2.1, and 𝑘 = 0.88 (giving the most apical frequency of 20 Hz).

In evaluating anatomical frequency mismatch with CIs, Greenwood’s equation lacks accuracy in at least one detail. It relates frequency to basilar membrane locations within the Organ of Corti (OC). Although CI electrodes are inserted into the cochlea, their target is the auditory nerve. More specifically, surviving spiral ganglion (SG) cell bodies located in Rosenthal’s canal within the modiolus are thought to be the primary target for CI stimulation rather than the radial nerve fibers that project from the SG into the OC which tend to degenerate (Stakhovskaya et al., 2007). Both the SG and OC spiral together along the modiolar axis, but differ in several respects. The SG length is shorter than the OC length by about 40%, the SG has a lower number of rotations about the modiolar axis (about 2 turns, OC has about 2 and 3/4 turns), and radial fibers do not follow a true radial (i.e., perpendicular) path from SG to OC instead becoming more skewed as one progresses towards the apex. Hence, Greenwood frequency-place estimates for the OC are not equivalent to those for the SG, even when SG and OC positions are normalized to their respective lengths, or angles of rotation. A useful anatomical correction was thus provided by Stakhovskaya et al. (2007). Analyzing human temporal bones, radial fibers were traced from OC to their respective locations along the SG, producing a mathematical function relating frequency-matched coordinates between OC and SG. More recently, Li et al. (2021) used synchrotron radiation phase-contrast imaging of human temporal bones to produce a 3-dimensional reconstruction of basilar membrane (BM) and SG positions, along with their radial nerve fiber projections. Applying Greenwood frequencies to BM positions and tracing individual dendrites from the BM to the SG at each frequency coordinate, they produced a “Greenwood-like” frequency-place function for the SG (i.e., Equation 1 with coefficient values that differed from those used to produce BM associated frequencies). The adjusted frequency-place functions for the SG provided by Stakhovskaya et al. (2007) and Li et al. (2021) differ slightly, particularly near the apex, but not by more than 0.5% in terms of SG position (as proportion of length) for any given frequency. Taken together, these adjusted SG frequency-place maps represent the current “gold-standard” for evaluating anatomical frequency mismatches in CIs. Importantly, both studies do not measure frequency dependance of OC (or BM) locations directly, relying instead on Greenwood’s function with commonly used coefficients.

However, relying on these coefficients as a standard for the human cochlear place-frequency map may be inaccurate in another detail. Known for some time, the frequency which drives maximal basilar membrane (BM) vibration at a particular cochlear location may also depend on input level. Some of the earliest, albeit indirect, evidence for this level dependence appear decades prior to Greenwood (1961) in studies of temporary threshold shift (TTS). In response to an intense fatiguing tone, the greatest threshold elevations occur at a frequency shifted by about 0.5 to 1 octave higher than that of the fatiguing tone (Perlman, 1941). This “half-octave” shift in TTS suggests that the BM traveling wave resulting from the intense fatiguing tone peaked at a more basal location, causing a threshold elevation at the frequency associated with that more basal location (McFadden, 1985; Johnstone et al., 1986). Another source of indirect evidence for a level dependent place-frequency shift in peak cochlear response comes from physiological measures of the cochlear microphonic (CM) in guinea pigs (Honrubia and Ward, 1968), showing large basalward shifts (4 mm) between points of maximum sensitivity to points of maximum voltage along the cochlear duct. However, this result has been questioned by Dallos (1973) who argues that the tendency of the nonlinear CM response with increasing sound level to produce greater saturation at apical relative to basal locations could produce such a shift even in the absence of any spatial shift of the underlying traveling wave.

Since these earlier studies, stronger evidence for level-dependent shifts in the cochlear place-frequency map have been observed directly from cochlear structures (BM, hair cells, supporting cells, auditory nerve fibers). For example, Russel and Nilsen (1997) measured BM displacement in vivo from guinea pigs at fifteen sites spanning 3 mm of cochlear distance in response to 15 kHz tones presented at various sound levels (15 to100 dB SPL). At lower sound levels, a narrow portion of the cochlea was active, peaking at the 15 kHz characteristic frequency (CF) place of around 14.5mm from the apex. At moderate sound levels up to about 60 dB SPL, the BM response broadened but retained a peak around the CF place. However, at levels greater than 60 dB SPL, the BM response continued broadening primarily at more basal locations, causing the peak response to shift basalward. The amount of shift in the peak BM response between lowest and highest sound levels was not large, about 0.5 mm, but demonstrates an example of basalward shift in the traveling wave peak with increasing level.

Studies like Russel and Nilsen (1997) are unique in the difficulty involved with successfully obtaining the spatial pattern of the traveling wave through many cochlear openings, which is highly traumatic to the cochlea. In contrast, a larger number of studies have obtained iso-intensity frequency responses at various input levels from cochlear structures at a fixed cochlear location through a single opening. If the traveling wave peak shifts basalward with increasing level, then the peak response at a fixed location would be expected to shift towards lower frequency inputs at higher stimulation levels. Two representative examples are presented in Figure 1, which shows iso-intensity frequency responses at various input levels recorded from an inner hair cell of a Mongolian gerbil (upper panel; Chaterjee and Zwislocki, 1997), and from an auditory nerve fiber of a squirrel monkey (lower panel; Rose et al., 1971). At low presentation levels (30 dB SPL) the peak response occurs at characteristic frequencies (CFs) of 2.4 kHz and 2.1 kHz (upper and lower panels, respectively), which persists up to around 50 dB SPL. At higher presentation levels, the frequency of the peak response (or best frequency, i.e. BF) shifts to lower frequencies. For example, at 80 dB SPL, the BF peak occurs at 1 kHz and 1.7 kHz (upper and lower panels, respectively). Considering the discrete and noisy nature of the measurements, one may also consider the BF corresponding to the centroid (instead of peak) of the 80 dB SPL iso-intensity contours which occur at 1.3 kHz and 1.5 kHz (upper and lower panels, respectively), i.e., one to one-half octave below CF (upper and lower panels, respectively). Although Figure 1 represents basalward shift with increasing level, the size and direction of these shifts are not necessarily consistent across cochlear locations, and across studies. For example, Geisler et al. (1974) and Dallos (1985) provide examples of iso-intensity responses showing relatively little shift. Furthermore, Rose et al. (1971) found the peculiar result that whereas nerve fibers with CFs greater than 1 kHz tended to produce BFs that shifted to lower frequencies at higher sound levels (i.e., basalward shift), those with CFs below 1 kHz tended to produce BFs that shifted to higher frequencies at higher sound levels.

**Figure 1:**
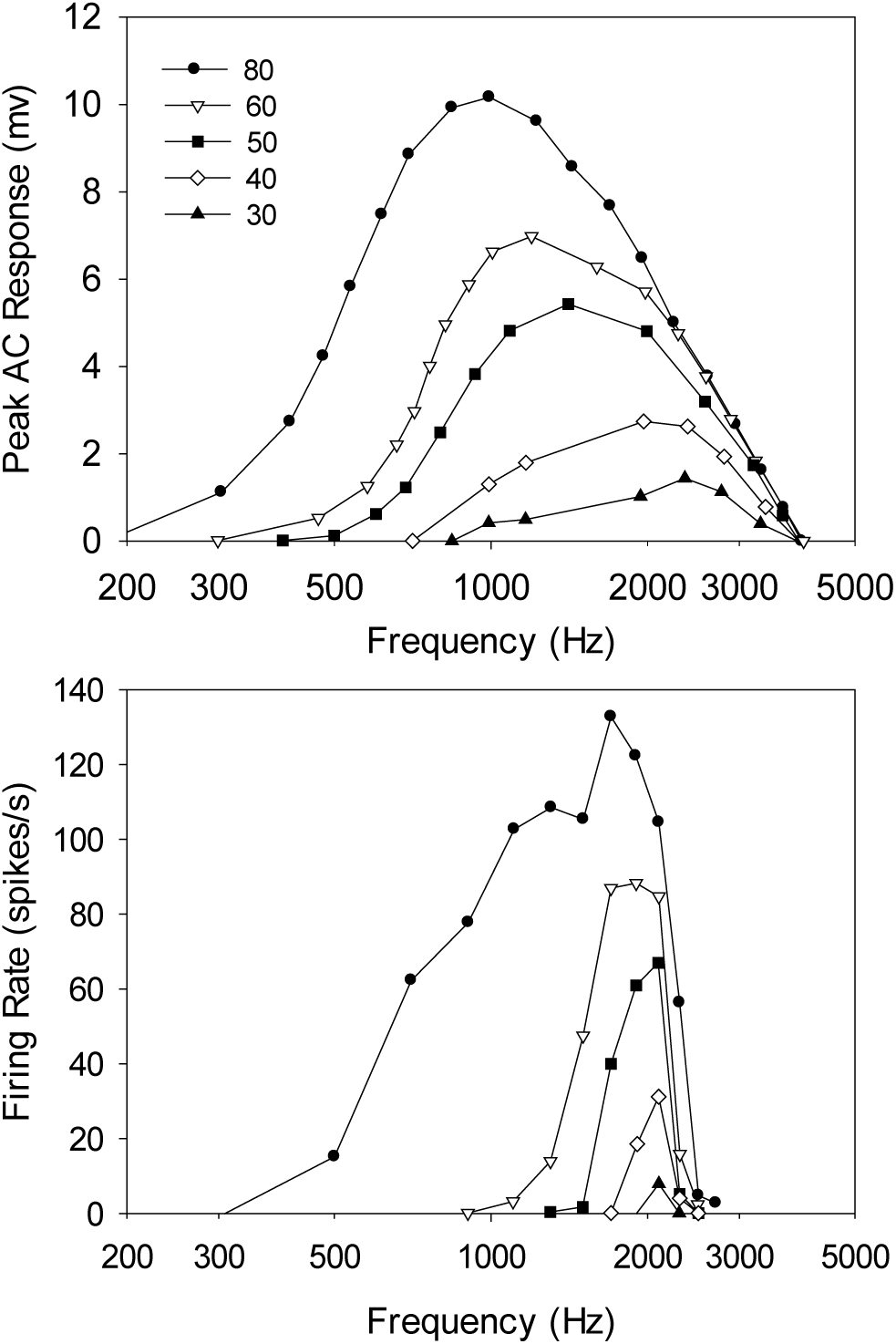
Iso-intensity contours at various input levels recorded from an inner hair cell of a Mongolian gerbil (upper panel; Chaterjee and Zwislocki, 1997), and from an auditory nerve fiber of a squirrel monkey (lower panel; Rose et al., 1971). At higher stimulation levels, best frequency (i.e., peak or centroid of frequency response) shifts away from characteristic frequency (i.e., peak frequency response at lowest input level). Downward shift in best frequency with increasing level is consistent with basalward shift of the traveling wave with increasing level.

A level-dependent shift in best frequency (BF) relative to characteristic frequency (CF) may have implications for evaluating anatomical frequency mismatch in CIs. For example, a basalward shift at higher presentation levels would reduce this mismatch. The purpose of the present study is to revisit Greenwood’s frequency-to-place map for humans in the context of level-dependent shifts in best-frequency (BF) relative to characteristic frequency (CF), to estimate the magnitude and direction of this shift using available physiological data from the literature, and to evaluate its impact on estimating anatomical frequency mismatch in CIs. We hypothesize that a BF-based place-frequency function may represent a more accurate tonotopic benchmark for CIs in comparison to current approaches based on CF alone (Stakhovskaya et al., 2007; Li et al., 2021).

## METHODS

### Source Materials

For the present study, place-frequency measurements were compiled from 15 physiological studies (Table 1). These studies include iso-intensity responses, like those of Figure 1, recorded at fixed locations (single-point) from in-vivo cochlear structures to tone stimuli at various frequencies and stimulation levels (e.g., Figure 1). Cochlear structures included in the present study were basilar membrane, Henson cells, inner and outer hair cells, and primary auditory nerve fibers. Data from each iso-intensity figure were digitized. The digitized data were used to evaluate characteristic frequency (CF) and best frequency most closely associated with a 70 dB SPL input (BF70). This level was chosen as it approximates moderately loud sounds experienced daily by normal hearing listeners, e.g., during conversation. CF was measured as the frequency of the peak response at the lowest input level obtained. BF70 was measured as the frequency associated with the centroid of the iso-intensity curve obtained at 70 dB SPL. For studies without an iso-intensity curve specifically at 70 dB SPL, BF70 values were interpolated from centroids of iso-intensity curves at the next higher and lower levels.

**Table 1:** Physiological studies used as source material for measurements of characteristic and best frequencies.

| # | Study | Figure(s) | Animal | Structure | Measurement Type |
| --- | --- | --- | --- | --- | --- |
| 1 | Chatterjee and Zwislocki (1997) | 3a, 4a-b | Mongolian Gerbil | IHC, Hensen Cell | AC response (mid and apical turn) |
| 2 | Dallos (1985) | 10a, 11a | Albino Guinea Pig | IHC, OHC | AC response (3<sup>rd</sup> turn) |
| 3 | Dallos (1986) | 3, 4 | Guinea Pig | IHC, OHC | AC response (3<sup>rd</sup> and 4<sup>th</sup> turn) |
| 4 | Geisler et al. (1974) | 2a-d, 7a-d | Squirrel Monkey | AN | Single unit discharge rate (basal region) |
| 5 | Muller and Robertson (1991) | 6a-b | Guinea Pig | AN | Single unit discharge rate |
| 6 | Overstreet et al. (2002) | 4a, 5a | Mongolian Gerbil | BM | BM vibration (approx. 1.2mm from base) |
| 7 | Recio-Spinosa and Oghalai (2017) | 2f-i | Albino Guinea Pig | BM | BM vibration (2 to 3.5 turns from base) |
| 8 | Recio-Spinosa and Oghalai (2018) | 1c, 1d | Albino Guinea Pig | BM | BM vibration (2 to 3.5 turns from base) |
| 9 | Ren and Nuttall (2001) | 2a | Mongolian Gerbil | BM | BM vibration (1<sup>st</sup> turn) |
| 10 | Rhode (2007) | 1c, 2c | Chinchilla | BM | BM vibration (3.5 - 4.25 mm from base) |
| 11 | Rhode and Recio (2000) | 3b, 3e, 3h, 11a | Chinchilla | BM | BM vibration (within 4mm of base) |
| 12 | Rose et al. (1971) | 1a-d, 2a-d, 4b | Squirrel Monkey | AN | Single unit discharge rate |
| 13 | Ruggero and Rich (1991) | 3 | Chinchilla | BM | BM vibration (3.5 mm from RW) |
| 14 | Ruggero et al. (1997) | 8, 10 | Chinchilla | BM | BM vibration (3.5 mm from OW) |
| 15 | Ruggero et al. (2000) | 1a | Chinchilla | BM | BM vibration (3.5 mm from OW) |

### Projecting CF and BF70 values from animals to humans

A simple relationship that follows from Greenwood’s general equation was used to project CF and BF70 measures obtained from the animal studies in Table 1 into estimates of equivalent human values. This relationship states that the ratio of frequencies associated with the same cochlear position in two mammalian species is constant throughout the length of the cochlea, equaling the ratio of each species’ upper frequency limit (i.e. most basal frequency). At the core of this relationship is an observation by Greenwood (1991) that longitudinal cochlear position relative to cochlear length is scale-related across species. That is, when cochlear position is expressed as a proportion of cochlear length, the Greenwood parameter “a” was found to be constant across species (i.e., a = 2.1). Greenwood also found that the parameter k varied only slightly across species (e.g. 0.8 ≤ k ≤ 1). If we assume that both of these parameters are held constant across two mammalian species being compared (e.g. a = 2.1 and k = 0.88), it follows

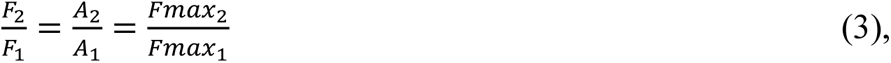

where F_1_ and F_2_ represent the frequencies associated with the same cochlear location (expressed as proportion of cochlear length) in the two species (subscripts 1 and 2); A_1_ and A_2_ represent the Greenwood parameters associated with the two species’ cochlear place-frequency map; and Fmax_1_ and Fmax_2_ represent the most basal frequencies in the two species.

For example, the CF associated with the half-way point along the cochlea in a cat is nearly 5 kHz (Liberman, 1982). The ratio in upper frequency limits of humans to cats is approximately 1/3 (e.g. 20 kHz in humans vs 60 kHz in cats), so the CF attributable to the half-way point along the cochlea in a human is about 1.67 kHz. This calculation becomes useful as there is, as yet, no direct way to carefully evaluate CF versus place for human cochlea in-vivo, whereas this has been done in other animal species. Equation 3 can also be applied to map BF70 values across species. For example, if a CF of 5 kHz shifts to a BF70 of 3.5 kHz at relatively high stimulation levels half-way along the cat cochlea, then the expected shift at a similar location in the human would be from a CF of 1.67 kHz to a BF70 of 1.17 kHz. In both cases, the frequency ratio from cat to human is 3:1, and the ratio of BF70 to CF is the same in each species (about 1:1.4 in this example).

Hence, CF and BF70 values measured from the studies in Table 1 were projected to human equivalent values by applying the ratio of each species upper frequency limit to that of humans. The upper frequency limit for a given species is subject to interpretation, and can vary across studies (Fay, 1988). Notwithstanding this caveat, the upper frequency limits chosen for the present study were as follows: 20 kHz for Humans, 57 kHz for cats, 60 kHz for Mongolian Gerbil, 50 kHz for Guinea Pig, 23 kHz for Chinchilla, and 46 kHz for Squirrel Monkey.

### Placement of CF and BF70 values onto Human Cochlear Place-Frequency Map

With CF and BF70 values projected from animals onto humans, the next step would be to assign these frequencies to their respective human position values along the cochlea. Traditionally, this has been done using Greenwood’s function with human-based coefficients of A = 165.4, a = 2.1, and k = 1 or 0.88 (depending on whether the most apical position is assigned 0 or 20 Hz, respectively). However, these coefficient values were provided irrespective of any shifts in cochlear place due to stimulation level. That is, Greenwood (1961, 1990) is equivocal to any differences that may occur between CF and BF at the same cochlear location. Hence, it is unclear from Greenwood alone whether it is more appropriate to apply the human-based coefficients to CF values in order to determine cochlear place while assuming these same place values for BF values, or whether to do the opposite (i.e. apply the human-based coefficients to BF values, etc.).

An alternative is to consider a cochlear place-frequency map from another species where CF as a function of cochlear position was measured directly, and use Equation (3) to transform this map into one for humans. Liberman (1982) constructed such a map from cat auditory nerve fibers. Characteristic frequencies were measured with threshold tuning curves, and the respective position of AN fibers along the Organ of Corti were measured using labeling techniques. The resulting place-frequency map closely followed a Greenwood relationship with coefficients A = 456, k = 0.8, and a = 2.1 (when cochlear length is expressed as a proportion, and frequency expressed in Hz). Transforming this map to humans with Equation (3) requires common values of coefficients k and a. The coefficient a = 2.1 is already the same in cats and humans. If the coefficient k is adjusted in cat so that k = 0.88 instead of k = 0.8, then the coefficient A for cat can be refit, changing slightly from A = 456 to A = 456.7. Assuming upper frequency values of 20 kHz in humans and 57 kHz in cats and applying Equation (1), the corresponding coefficient A for human cochlear place vs CF becomes A = 160. This derived value for A is very close to A = 165.4 (indeed the latter would be obtained if we chose an upper frequency limit of 20.67 kHz instead of 20 kHz). This outcome suggests that Greenwood’s equation, with his human-based coefficients, provides a good description of the human place-frequency map associated with CF, but (to the extent that the two differ) a less accurate representation of the human place-frequency map associated with BF.

For simplicity, and consistency with its usage in previous studies, we adopt values of A = 165.4, k = 2.1, and a = 0.88 to represent human cochlear place values associated with CF (and not BF at higher levels), relying on our use of Equation (3) and Liberman (1982) as justification. Cochlear position values for BF values are assumed equal to those obtained for CF values via Greenwood, since CF and BF values from the studies in Table 1 were obtained from the same cochlear location.

### Placement of CF and BF values onto Human Spiral Ganglion map

Our application of Greenwood to CF values provides cochlear locations in units of proportion of cochlear length starting from the apex. To compare CF and, by extension, BF locations more easily to frequencies mapped to cochlear implant stimulation sites, we are ultimately interested in transforming CF and BF locations to associated spiral ganglion locations in units of angle of rotation. Based on anatomical measures from human temporal bone, Stakhovskaya et al. (2007) provide these transformations, but express cochlear and spiral ganglion locations in units of percent length from the base instead of apex. Hence, we first convert our Greenwood derived locations from proportion of cochlear length from the apex to a percentage of cochlear length from the base (i.e., subtract from 1 and multiply by 100). Percentage of cochlear length from the base (more specifically, organ of corti length or OC%) can then be transformed to percent distance of the corresponding location along the spiral ganglion (SG%) using the following relationship (i.e., Equation 1 in Stakhovskaya et al., 2007):

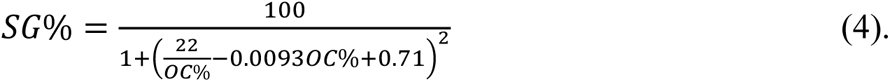

Percent distance of spiral ganglion from the base (SG%) can then be transformed to angle of rotation (SG°, in degrees) using the following relationship (i.e., inverting Equation 4 in Stakhovskaya et al., 2007):

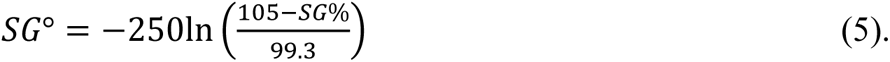

When deriving their frequency map for spiral ganglion location, Stakhovskaya et al. (2007) traced the spiral ganglion locations to their corresponding organ of corti locations, and then assigned frequencies to the latter using Greenwood’s equation with the same coefficients used in the present study (A = 165.4, k = 2.1, and a = 0.88). Considering Lieberman’s (1982) place-frequency map for the cat, and the ratio of upper frequency limits of hearing in cat vs human, we suggest that these Greenwood coefficients provide a human place-frequency map most closely associated with CF (as opposed BF). Hence, we used Equations 4 and 5 to evaluate a spiral ganglion angle of rotation for each CF, and then apply the same location to associated BF values. That is, we chose to normalize our CF values to the spiral ganglion place-frequency map provided by Stakhovskaya et al. (2007), in order to evaluate the extent of shift in associated BF values. BF values are then compared to frequencies typically associated with cochlear implant stimulation sites using values provided by Landsberger et al., 2015.

## RESULTS

The shift between characteristic frequencies (CF) and centroid-based best frequencies at 70 dB SPL (BF70) as a function of spiral ganglion locations are depicted in Table 2 and in Figure 2. In Table 2, CF and BF70 values extracted from the animal studies in Table 1 are listed next to their respective study number and figure from which data were digitized. These animal-based CF and BF70 values were projected to human CF and BF70 values by applying the appropriate human-to-animal upper frequency limit ratio (Equation 3 of the present study). Spiral ganglion locations (in degrees of rotation from base) for projected human CF values were derived using Equation 2 of the present study with A = 165.4, k = 2.1, and a = 0.88, and then applying Equations 4 and 5 (i.e., Stakhovskaya et al., 2007). CF spiral ganglion locations were also applied to corresponding BF70 values because both were obtained at the same cochlear location. The data in Table 2 are sorted by increasing values of human CF to assist with interpreting shifts between BF70 and CF at different cochlear locations. Overall, the difference between CF and BF70 human estimates was statistically significant (signed-rank test z = 5.143, p < 0.001), but the amount and direction of shift depended on cochlear location. In Table 2, the direction and degree of this shift is characterized by the ratio of BF70 to CF (which is assumed to be the same for animal and human measures). Ratio values less than one (i.e., BF70 < CF) indicate a basalward shift in best-frequency locations relative to characteristic frequency locations. For example, if BF70 was a half-octave lower than CF at a cochlear location with CF = 2 kHz, then BF70 = 1,414 Hz at this location and BF70/CF = 0.71. This represents a basalward shift because one must move more basally from this location to find the BF70 value equal to 2 kHz. Conversely, ratio values greater than one (i.e., BF70 > CF) indicate an apicalward shift in best-frequency locations relative to characteristic frequency locations.

**Figure 2:**
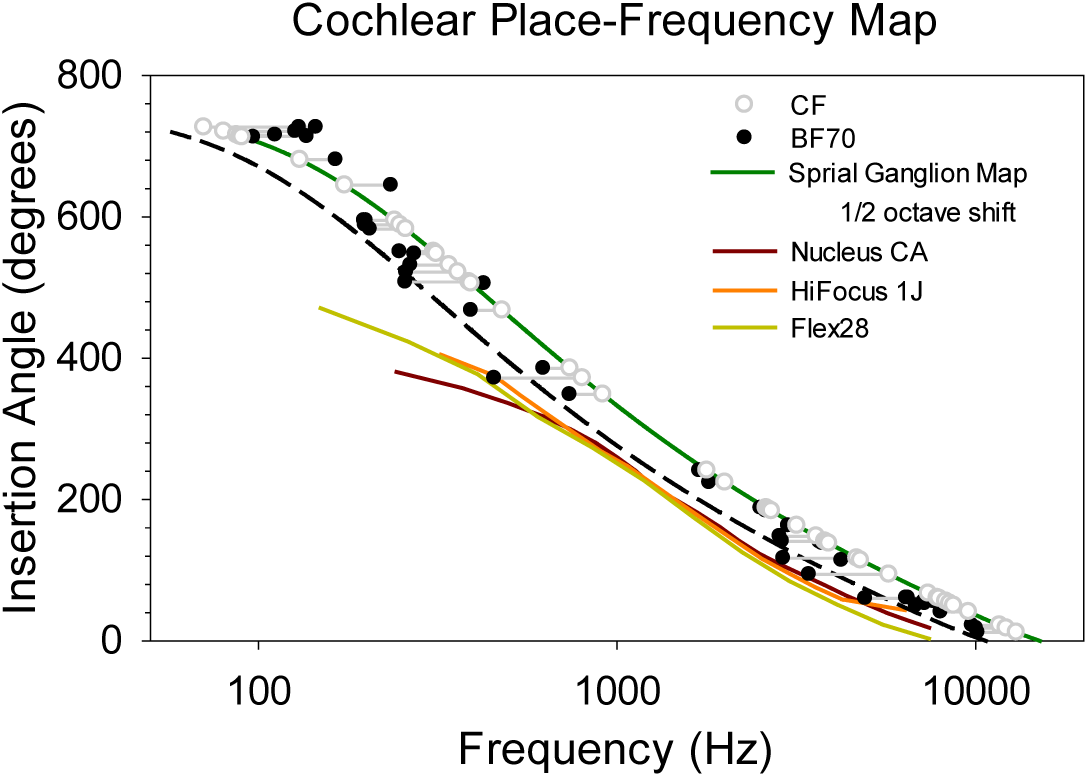
Shift in human projected CF and BF70 values (unfilled gray and filled black circles, respectively) as a function of spiral ganglion locations. CF values are overlayed onto the spiral ganglion place-frequency map (green line) derived from Stakhovskaya et al. (2007). Spiral ganglion locations of electrodes for three cochlear implant devices and their typically assigned center frequencies shown in red, orange, and yellow (Landsberger et al., 2015). The dashed line represents the spiral ganglion place-frequency map shifted downwards by a half-octave.

**Table 2:** Values of characteristic frequency (CF) and best frequency centroids for 70 dB SPL iso-intensity contours (BF70) extracted from animal studies in Table 1 listing associated figures. CF and BF70 values scaled to human values based on human to animal upper frequency ratios. Human CF values transformed to spiral ganglion angular position from base. Table sorted by increasing human CF values for easier reference to Fig 2.

| Study # | Fig. | Animal (kHz) |  | Human (kHz) |  |  | BF70/CF |
| --- | --- | --- | --- | --- | --- | --- | --- |
|  |  | CF | BF70 | SG° | CF | BF70 |  |
| 8 | 1c | 0.176 | 0.324 | 727 | 0.070 | 0.129 | 1.84 |
| 8 | 1d | 0.176 | 0.362 | 727 | 0.070 | 0.144 | 2.06 |
| 7 | 2f | 0.200 | 0.317 | 721 | 0.080 | 0.127 | 1.59 |
| 12 | 1a | 0.200 | 0.256 | 716 | 0.087 | 0.111 | 1.28 |
| 7 | 2g | 0.222 | 0.341 | 714 | 0.089 | 0.136 | 1.53 |
| 3 | 3 left | 0.225 | 0.242 | 713 | 0.090 | 0.097 | 1.08 |
| 12 | 1b | 0.300 | 0.378 | 681 | 0.130 | 0.164 | 1.26 |
| 12 | 1c | 0.400 | 0.539 | 645 | 0.174 | 0.234 | 1.34 |
| 7 | 2h | 0.600 | 0.498 | 595 | 0.240 | 0.199 | 0.83 |
| 7 | 2i | 0.600 | 0.492 | 595 | 0.240 | 0.197 | 0.82 |
| 1 | 4b | 0.744 | 0.594 | 589 | 0.248 | 0.198 | 0.80 |
| 1 | 4a | 0.773 | 0.614 | 583 | 0.258 | 0.205 | 0.79 |
| 3 | 4 left | 0.770 | 0.618 | 551 | 0.308 | 0.247 | 0.80 |
| 2 | 11a | 0.783 | 0.679 | 548 | 0.313 | 0.272 | 0.87 |
| 2 | 10a | 0.850 | 0.663 | 532 | 0.340 | 0.265 | 0.78 |
| 3 | 3 right | 0.900 | 0.645 | 522 | 0.360 | 0.258 | 0.72 |
| 3 | 4 right | 0.966 | 0.642 | 508 | 0.386 | 0.257 | 0.67 |
| 12 | 4b | 0.900 | 0.978 | 506 | 0.391 | 0.425 | 1.09 |
| 12 | 1d | 1.100 | 0.900 | 468 | 0.478 | 0.391 | 0.82 |
| 12 | 2a | 1.700 | 1.432 | 386 | 0.739 | 0.623 | 0.84 |
| 1 | 3a | 2.400 | 1.362 | 372 | 0.800 | 0.454 | 0.57 |
| 12 | 2b | 2.100 | 1.695 | 349 | 0.913 | 0.737 | 0.81 |
| 12 | 2c | 4.100 | 3.900 | 241 | 1.783 | 1.696 | 0.95 |
| 4 | 2a | 4.603 | 4.158 | 224 | 2.001 | 1.808 | 0.90 |
| 4 | 7b | 6.000 | 5.781 | 188 | 2.609 | 2.513 | 0.96 |
| 4 | 7c | 6.100 | 5.944 | 186 | 2.652 | 2.584 | 0.97 |
| 4 | 7a | 6.200 | 5.972 | 184 | 2.696 | 2.597 | 0.96 |
| 4 | 2b | 7.305 | 6.900 | 163 | 3.176 | 3.000 | 0.94 |
| 5 | 6a | 9.000 | 7.103 | 148 | 3.600 | 2.841 | 0.79 |
| 5 | 6b | 9.500 | 7.214 | 141 | 3.800 | 2.886 | 0.76 |
| 4 | 7d | 8.807 | 8.442 | 140 | 3.829 | 3.670 | 0.96 |
| 4 | 2d | 8.966 | 8.578 | 138 | 3.898 | 3.730 | 0.96 |
| 9 | 2a | 14.00 | 8.730 | 117 | 4.667 | 2.910 | 0.62 |
| 11 | 3b | 5.500 | 4.860 | 114 | 4.783 | 4.226 | 0.88 |
| 10 | 1c | 6.600 | 3.947 | 94 | 5.739 | 3.432 | 0.60 |
| 12 | 2d | 17.000 | 17.07 | 67 | 7.391 | 7.423 | 1.00 |
| 13 | 3 | 9.000 | 7.454 | 61 | 7.826 | 6.482 | 0.83 |
| 14 | 10 | 9.000 | 7.360 | 61 | 7.826 | 6.400 | 0.82 |
| 10 | 2c | 9.100 | 5.659 | 60 | 7.913 | 4.921 | 0.62 |
| 4 | 2c | 19.00 | 17.50 | 56 | 8.261 | 7.609 | 0.92 |
| 15 | 1a | 9.500 | 8.262 | 56 | 8.261 | 7.184 | 0.87 |
| 11 | 3e | 9.750 | 8.293 | 53 | 8.478 | 7.211 | 0.85 |
| 14 | 8 | 10.00 | 7.841 | 50 | 8.696 | 6.819 | 0.78 |
| 11 | 11a | 11.00 | 9.187 | 41 | 9.565 | 7.989 | 0.84 |
| 6 | 4a | 35.00 | 29.30 | 22 | 11.67 | 9.767 | 0.84 |
| 11 | 3h | 14.00 | 11.52 | 18 | 12.17 | 10.02 | 0.82 |
| 6 | 5a | 39.00 | 30.40 | 12 | 13.00 | 10.13 | 0.78 |

Nearly three-quarters of the CF and BF70 measurements obtained had BF70 values that were shifted away from CF by more than 10% (i.e., BF70/CF < 0.9 or BF70/CF > 1.1). For spiral ganglion locations up to 600° from the base, the shift tended to be basalward with BF70 values shifting downward in frequency by an average of almost a third-octave down relative to CF values at the same location (i.e. mean BF70/CF = 0.83). For spiral ganglion locations greater than 600° from the base, shifts became more apicalward (i.e., BF70/CF > 1), with BF70 values shifting upward in frequency by an average of just over a half-octave relative to CF values at the same location (i.e., BF70/CF = 1.5). A spiral ganglion location of 600° from the base corresponds to a CF value of about 230 Hz in humans.

In Figure 2, CF values (unfilled gray circles) are overlayed onto the spiral ganglion place-frequency map (green line) derived from Stakhovskaya et al. (2007). For spiral ganglion locations less than 600° from the base, BF70 values (filled black circles) tend to lie to the left of spiral ganglion place-frequency map, many of which cluster near the black dashed line representing the spiral ganglion map down-shifted in frequency by a half-octave. To serve as a reference, Figure 2 includes average cochlear locations of implanted electrodes for specific devices from three cochlear implant manufacturers as a function of the typical center frequencies assigned to those electrodes (red, orange, and yellow lines; data from Landsberger et al., 2015). The prevailing view is that cochlear implant electrodes stimulate neurons in the spiral ganglia, as opposed to the radial fibers that extend into the cochlea which tend to degenerate in severe to profound hearing loss (Stakhovskaya et al., 2007). Hence, comparing the proximity of electrode place-frequency locations to spiral ganglion place-frequency locations provides a measure of tonotopic mismatch that may be experienced by CI users. When considering the spiral ganglion map associated with CF values, this tonotopic mismatch is approximately one octave or more. However, when considering spiral ganglion locations associated with BF70 values, the tonotopic mismatch is less extreme, particularly for middle and basal electrodes. Nevertheless, a relatively large mismatch persists for more apical electrodes.

Considering the BF70/CF values in Table 2, an approximate approach to adjusting tonotopic frequencies associated with cochlear implant electrode locations for a moderately loud presentation level would be to downshift spiral ganglion characteristic frequencies by a factor of 0.8 (i.e., 1/3 octave or 4 semitones) for electrode locations up to 600° from the base, and upshift CF values by a factor of 1.4 (i.e., 1/2 octave or 6 semitones) for more apical electrode locations beyond 600°. For a more numerical approach, non-linear regression was used to determine a best-fit to the BF70/CF data as a function of spiral ganglion location as shown in Figure 3. When spiral ganglion location is measured as percent length from the base (i.e., SG% as in Equation 4), the following empirical relationship produced an R^2^ = 0.77:

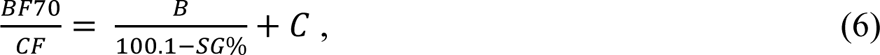

**Figure 3:**
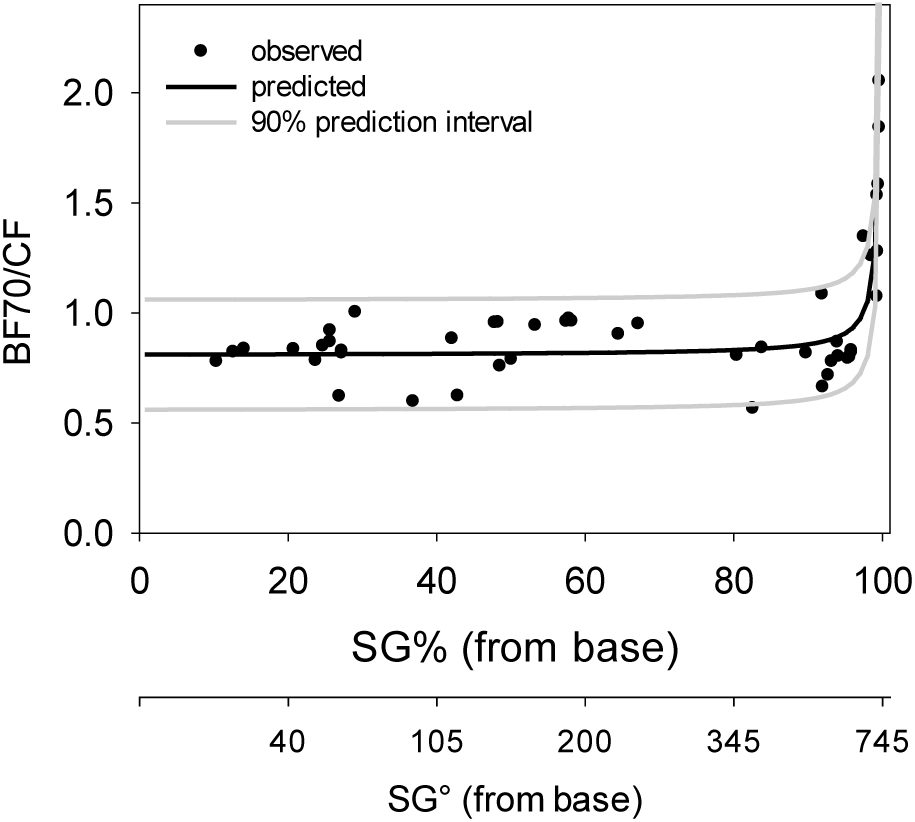
Nonlinear regression (R^2^ = 0.77) of Equation 6 to BF70/CF data from Table 2 as a function of spiral ganglion location.

with mean coefficient values (and their 95% confidence interval) of B = 0.523 ([0.438, 0.609]), and C = 0.806 ([0.757, 0.855]). The nonlinear regression in Figure 3 also includes 90% prediction intervals for the BF70/CF data based on Equation 6.

Combining Equations 5 and 6, one can produce BF70/CF ratios at any desired spiral ganglion location (in degrees from the base). These ratios can serve as a level-dependent correction factor to apply to spiral ganglion CF values at each respective location. For example, at 360° from the base, the spiral ganglion CF value is 856 Hz (Equations 1, 4, and 5 with A = 165.4, k = 2.1, and a = 0.88). At this location, mean BF70/CF = 0.834 (Equation 6) with a 90% prediction interval of [0.584, 1.084]. Multiplying the spiral ganglion CF at 360° from the base by these ratio values gives a mean BF70 = 714 Hz with a 90% prediction interval of [500 Hz, 928 Hz]. Applying this approach, CF and mean BF70 values (along with upper and lower 90% prediction intervals) are shown in Table 3 at spiral ganglion locations from 0° through 720° from the base in 45° increments (a similar Table in 1° increments is available online as supplementary material).

**Table 3:**
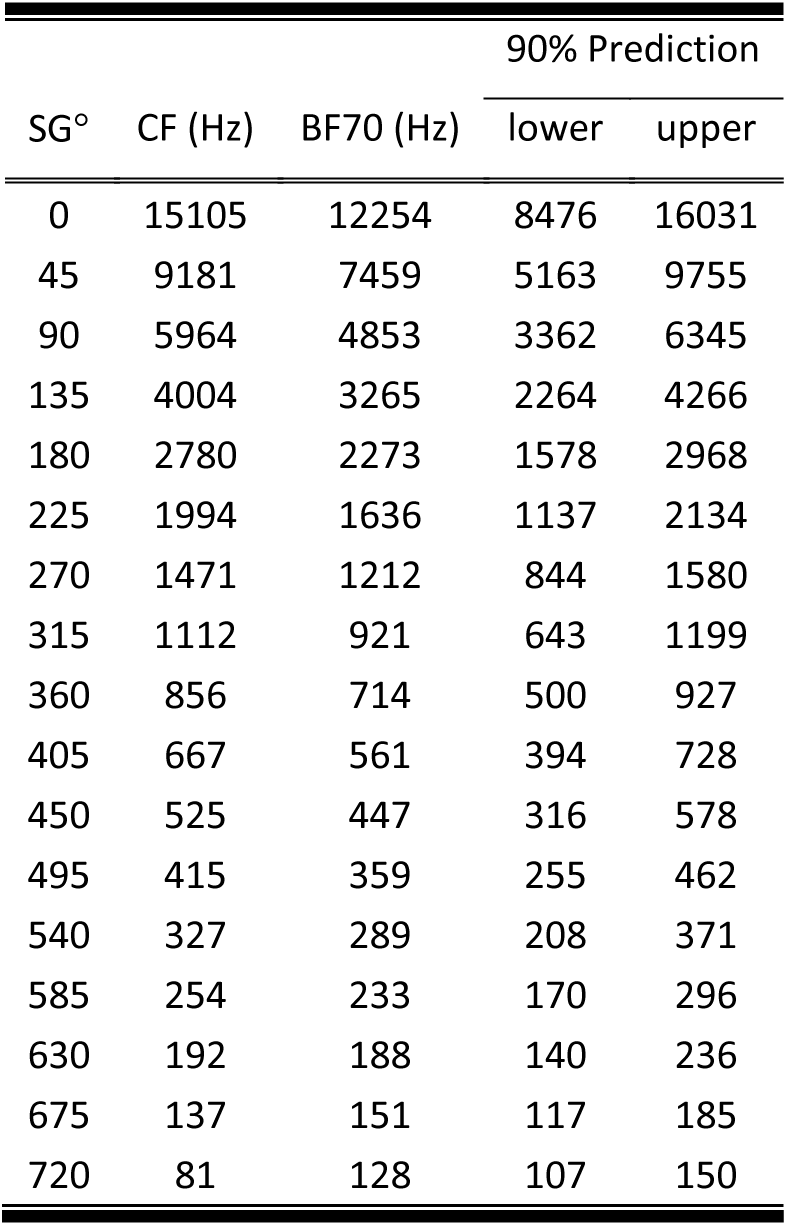
Mean level-dependent correction (BF70) of characteristic frequency (CF) values as a function of spiral ganglion location (in degrees), along with lower and upper 90% prediction interval for BF70.

| SG° | CF (Hz) | BF70 (Hz) | 90% Prediction |  |
| --- | --- | --- | --- | --- |
|  |  |  | lower | upper |
| 0 | 15105 | 12254 | 8476 | 16031 |
| 45 | 9181 | 7459 | 5163 | 9755 |
| 90 | 5964 | 4853 | 3362 | 6345 |
| 135 | 4004 | 3265 | 2264 | 4266 |
| 180 | 2780 | 2273 | 1578 | 2968 |
| 225 | 1994 | 1636 | 1137 | 2134 |
| 270 | 1471 | 1212 | 844 | 1580 |
| 315 | 1112 | 921 | 643 | 1199 |
| 360 | 856 | 714 | 500 | 927 |
| 405 | 667 | 561 | 394 | 728 |
| 450 | 525 | 447 | 316 | 578 |
| 495 | 415 | 359 | 255 | 462 |
| 540 | 327 | 289 | 208 | 371 |
| 585 | 254 | 233 | 170 | 296 |
| 630 | 192 | 188 | 140 | 236 |
| 675 | 137 | 151 | 117 | 185 |
| 720 | 81 | 128 | 107 | 150 |

## DISCUSSION

### Implications for Cochlear Implants

The present study demonstrates that the cochlear place-frequency map is level dependent, and a correction is offered for the difference between a spiral-ganglion (SG) place-frequency map at characteristic frequency (CF), and one at 70 dB SPL (BF70). A level of 70 dB SPL was chosen as it is representative of a moderately loud conversational level in listeners with normal hearing (NH), and may reflect a more realistic tonotopic reference when considering anatomical frequency mismatch inherent in cochlear implant (CI) standard-of-care frequency-to-electrode assignments. Figure 4 shows the average angle of insertion of individual electrodes and their standard-of-care frequency allocation for the three CI devices plotted in Figure 2 (from Landsberger et al., 2015), as well as the SG place-frequency map at CF (green dashed line) and at BF70 (solid black line), along with the latter’s upper and lower 90% prediction interval (solid gray lines). The amount of place-frequency mismatch using default frequency allocations is less extreme when BF70, instead of CF, is considered as the tonotopic reference. Indeed, most electrodes straddle the lower 90% prediction interval for BF70, which suggests the possibility of anatomical frequency mismatch being significant for only the most apical electrodes. On the other hand, the upper 90% prediction interval straddles CF, supporting the previously held notion that average frequency mismatch is an octave or greater for typical insertions in these devices. On average, the present study suggests that anatomical frequency mismatch is about 4 semitones lower when using BF70, instead of CF, as the tonotopic reference. Hence, although the present study does not entirely refute the previously held notion of large anatomical frequency mismatch in standard-of-care fittings, it does suggest that the mismatch is likely to be less extreme and possibly non-existent for all but the most apical electrodes.

**Figure 4:**
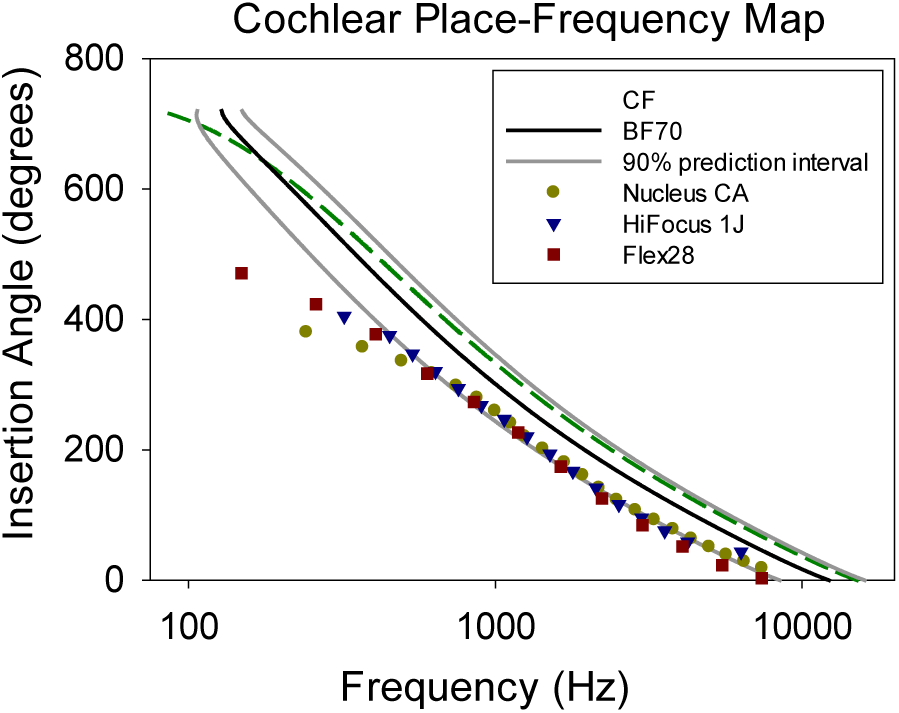
Average angle of insertion of individual electrodes and their standard-of-care frequency allocation for the three CI devices (symbols, from Landsberger et al., 2015), as well as the spiral ganglion place-frequency map at CF (green dashed line) and its level-dependent correction, i.e., BF70 (solid black line), along with the latter’s upper and lower 90% prediction interval (solid gray lines).

One implication of these findings is that correcting for anatomical frequency mismatch in CIs may require a less extreme change to standard-of-care frequency allocations than previously thought. Whether this change will amount to perceived benefit over standard-of-care settings is unclear. Although several studies have demonstrated varying degrees of benefit when using anatomical-based fitting (Di Maro et al., 2022; Dillon et al., 2023; Fan et al., 2023; Kurtz et al., 2023; Tóth et al., 2023; Creff et al., 2024; Dessard et al., 2024), others have demonstrated either no benefit or poorer performance in comparison to standard-of-care (Grasmeder et al., 2014; Jiam et al., 2018; Dirks et al., 2022; Lambriks et al., 2023). Part of the problem is in how the anatomical-based place-frequency map is defined. When assigning frequencies to electrodes, some studies apply Greenwood’s human coefficients (with and/or without spiral ganglion adjustments) more strictly across all electrodes (e.g. Grasmeder et al., 2014), whereas other studies limit the application of Greenwood to mid-frequency electrodes while anchoring the more extreme apical and/or basal electrodes to standard-of-care frequencies (e.g. Di Maro et al., 2022). On one hand, stricter application of Greenwood frequencies to all electrodes can result in greater degradation of speech cues, as it may involve removal of substantial low-frequency information while compressing the remaining frequency range to a narrower cochlear region (i.e., after deactivation of basal electrodes with anatomically-based frequencies that exceed the upper limit of the CI system). On the other hand, the cost of reduced speech information may be outweighed by the benefit of an anatomically-based frequency map that reduces frequency mismatch, e.g. for subjects who don’t fully adapt to the standard of care, or for subjects where an anatomically-based frequency map provides better integration with available hearing in the contralateral ear (e.g. bimodal CI users and those with single-sided deafness). The latter argument assumes, of course, that Greenwood’s anatomically-based frequency map for humans is, in fact, the correct anatomical reference. In the present study, we argue for a less extreme anatomical reference for CI users, downshifted in frequency by about 4 semitones, on average, relative to spiral ganglion maps based on Greenwood’s human coefficients.

Coincidentally, Creff et al., 2024 showed speech understanding benefit when using non-standard vs conventional frequency maps aimed at reducing frequency mismatch that appear to closely match the level-corrected tonotopic reference of the present study (i.e., BF70). Tonotopic fitting was achieved via postoperative flat-panel CT for each subject by measuring the cochlear duct length from the round window to a given electrode and then using Greenwood’s (1990) human coefficients to determine the associated frequency. What’s unique about Creff et al. in comparison to similar studies that implement Greenwood, is that the Greenwood derived frequencies were assigned to the upper edge of the electrode’s frequency passband instead of its center frequency. This was done for electrode locations associated with frequencies up to 3 kHz, beyond which frequencies were distributed logarithmically so that the upper edge of the most basal electrode was assigned 8500 Hz. Assigning Greenwood frequencies to the upper edge of an electrode’s passband resulted in its center frequencies becoming shifted down by about 3 semitones, which is near the average level-corrected shift described in the present study. Although indirect, Creff et al. (2024) offers support for achieving speech understanding benefit when using a non-standard frequency allocation that reduces frequency mismatch, but that is less extreme than place-frequency maps derived more strictly from Greenwood’s (1990) human coefficients (and associated spiral ganglion corrections). The present study provides justification for why such an intermediary map may be, in fact, a more accurate tonotopic reference.

Of course, the above discussion applies to insertion angles up to about 600°. At deeper insertion angles, the level-corrected adjustment of the present study no longer manifests as a downward shift in frequency relative to the spiral ganglion CF place-frequency map, but rather, an upward frequency shift (Figure 4). This upward shift has implications for place coding in the cochlear apex. That is, at higher stimulation levels, the BF place-frequency map slopes upward in the apical region such that larger changes in place become associated with smaller changes in frequency, meaning an overall reduction in representation of place-based cues in the cochlear apex. The situation is more extreme for cochlear implants where stimulation occurs at the spiral ganglion. Even without the level-corrected adjustment of the present study, Li et al. (2021) noted a marked drop in place-frequency specificity in their spiral ganglion function beyond 600°. Taken together, there appears to be less benefit in delivering place-based frequency information to these more apical regions. This observation is consistent with Canfarotta et al. (2022) who found a small but significant positive correlation between insertion angle and word scores, but which plateaued beyond 600°.

### Qualifications and Limitations

In the present study, the SG place-frequency values at CF (Table 3) differ somewhat from those tabulated in Stakhovskaya et al. (2007), mostly undershooting the latter by about 5% to 10%, although we used the same Greenwood coefficients. The discrepancy appears to be due to deviations between the empirical equation reported by Stakhovskaya et al. relating spiral ganglion length to degrees of rotation (i.e., our source for Equation 5 of the present manuscript) and the actual means of the observed data tabulated in Stakhovskaya et al., 2007 (c.f. their Figure 9). Our SG place-frequency values at CF are also comparable to those reported by Li et al. (2021), mostly overshooting the latter by about 5%. Notwithstanding these differences, our CF to BF70 correction (Equation 6) can be applied to whichever variety of the SG place-frequency map at CF one chooses to adopt.

The correction provided in the present study is based on centroid measurements made from digitized iso-intensity responses at 70 dB SPL (BF70), culled from published figures spanning nearly 40 years. Without the original source data, it is not possible to gauge the degree of error inherent in the extraction of this data via digitization. One mitigating factor is that the most essential aspect of the data is the ratio of BF70 and CF, which may be less susceptible to errors of digitization in comparison to the absolute values of either variable. A further caveat regarding the BF70 correction reported herein is that the studies cited as source materials for the present study (Table 1) varied in the degree of surgical manipulation associated with stimulus delivery and recording of cochlear responses, which can alter response characteristics. For example, to access cochlear structures, some studies required opening the auditory bulla widely which can elevate low-frequency responses by as much as 16 dB (Ruggero et al., 1990). This effect diminishes with increasing frequency, becoming negligible at around 700 Hz in chinchilla (i.e., about 600 Hz in humans). Other examples of manipulations that could alter normal cochlear response include severing of middle ear muscles, loading the cochlea with reflective beads to aid visualization of basilar membrane vibration, and the degree to which the inner ear was disturbed to obtain cochlear responses (Narayan et al., 1998). These factors likely contributed to the variation in BF70 shifts observed in the present study.

Aside from more apical regions, the peak (or centroid) of the cochlear response to a 70 dB SPL tone tends to shift downwards in frequency relative to CF. This downward shift supports the observation that the peak (or centroid) of the traveling wave envelope moves to a more basal location at higher presentation levels (Russel and Nilsen, 1997). Higher intensity levels would result in further shift, likely approaching the limits predicted by the half-octave shift phenomenon, and perhaps beyond (McFadden, 1985; Johnstone et al., 1986). This shift is thought to be due to the interplay of active and passive mechanical properties of the cochlear partition (Davis, 1983). At lower levels, active mechanics predominate, and the cochlear response is amplified in a narrow region near the apical edge of the traveling wave envelope. At higher levels, passive mechanics predominate, and the traveling wave envelope broadens asymmetrically toward the base while remaining relatively anchored at its apical edge (Russel and Nilsen, 1997). The net result is a basalward shift in the traveling wave envelope peak (or centroid) with increasing level. In the more apical regions, the opposite effect appears. Namely, the peak cochlear response shifts to higher frequencies with increasing level suggesting more apicalward movement of the traveling wave. Indeed, Békésy (1947) confirmed as much in his human temporal bone preparations, noting that the traveling wave in response to 200 Hz at around 30 mm from the stapes becomes displaced toward the helicotrema with increasing amplitude. It is unclear, however, to what extent these interpretations of the traveling wave hold at the very extremes of the cochlea, where shifts in peak response frequency with level may instead reflect a more complicated interaction between the mechanical properties of the cochlea and its physical boundaries.

### Greenwood’s human coefficients and characteristic frequency

In the cochlear implant field, it has become customary to associate Greenwood’s (1961, 1990) human coefficients (𝐴 = 165.4, 𝑎 = 2.1, and 𝑘 = 0.88) with characteristic frequency (CF), despite limited physiological support in humans. To qualify, Greenwood (1961, 1990) provided abundant physiological support for the validity of his place-frequency function across several animal species, each one producing species-specific coefficients. Coefficients 𝑎 (when cochlear position is expressed as proportion of cochlear length) and 𝑘 remained relatively stable across species, and coefficient 𝐴 varied widely across species depending on their upper frequency limit of hearing. In the case of humans, by contrast, Greenwood’s coefficients were derived from fitting psychophysical critical bandwidth data to Greenwood’s function, which itself arose from the assumption that critical band number is proportional to distance, in mm, along the cochlea (Greenwood, 1961). Human physiological support was provided in the form of Békésy (1960) and Kringlebotn et al. (1978) who obtained post-mortem place-frequency measures in human temporal bones at very high stimulation levels (equivalent to ≥ 120 dB SPL). Although Greenwood (1990) provided justification for why these measures would not differ greatly from similar in-vivo measurements, it’s been known for a while that cochlear responses at high stimulation levels are not representative of responses at lower stimulation levels (first shown by Rhode, 1971), likely due to a non-linear interplay of passive and active cochlear mechanics (Davis, 1983). Furthermore, upon closer inspection, the Békésy (1960) data appear shifted to more basalward positions at a given frequency (or to lower frequencies at a fixed place) in comparison to Greenwood’s function with human coefficients. The data from Kringlebotn et al. (1978) do conform more closely to Greenwood’s function with human coefficients, but seem to be based on a local maximum in the cochlear frequency response rather than on the actual maximum (see Fig. 2 in Kringlebotn et al.). Had they used the actual peak of that function, the frequency associated with that location would have been about 1.5 octaves lower than the value provided in the paper (about 1 kHz instead of about 3.6 kHz), consistent with a basalward shift. Given these observations, interpreting Greenwood’s human coefficients as representing a place-frequency map associated with CF would be an inference based on indirect observation from limited data.

In the present study, Greenwood’s human coefficients were chosen as representative of CF, but not by relying on limited human physiological support. Instead, a simple animal-to-human conversion based on each species’ upper audible frequency limit (Equation 3) was applied to the comprehensive data of Liberman (1982) obtained in cats. This data represents the strongest evidence to date relating cochlear place to characteristic frequency, unique in the degree of care used to directly obtain CF values of auditory nerves in-vivo and trace their BM locations. Applying Equation 3 to Liberman (1982) yielded a projected value of 𝐴 = 160 in humans, which closely resembles the commonly used value of 𝐴 = 165.4. Hence, a reasonable transformation of the Liberman data to the human case shows that Greenwood’s human coefficients are representative of CF, confirming the implicit assumption made by Stakhovskaya et al. (2007) and Li et al. (2021), whose validity is inextricably tied to these coefficients.

## Conclusion

Anatomically based place-frequency functions used in the cochlear implant field rely on Greenwood’s (1961, 1990) human coefficients to relate cochlear place and characteristic frequency, even though there is a very limited amount of human physiological data supporting its validity. In addition, known for several decades, the frequency associated with the peak (or centroid) of the cochlear response (i.e., best frequency) tends to shift away from characteristic frequency at higher stimulation levels. After surveying a relatively large number of animal studies and implementing a simple animal-to-human transformation, the present study provides further confirmation for interpreting Greenwood’s human coefficients as relating cochlear place to characteristic frequency, and proposes a novel place-frequency function that, to our knowledge, is the first to estimate how much place-frequency varies with stimulation level. By correcting for the shift in cochlear place-frequency responses at a moderately loud conversational level of 70 dB SPL, the proposed function results in place-frequency corrections that are, on average, about 4 semitones smaller than those indicated by currently utilized place-frequency functions based on Greenwood’s human coefficients and associated spiral-ganglion corrections. This basalward shift occurs for insertion angles up to about 600° from the base, beyond which the shift reverses direction becoming apicalward. This level-dependent place frequency function has implications for frequency-to-electrode assignments in cochlear implants, suggesting that the anatomical frequency mismatch inherent in standard-of-care fittings may be less extreme than previously thought. Overall, it is argued that the level-dependent frequency-place function put forth in this study provides a more accurate tonotopic reference that could guide future approaches to optimizing frequency-to-electrode assignments in cochlear implant users.

## ACKNOWLEDGEMENTS

This research was supported by NIH/NIDCD grants DC020293 (PI: Sagi) and DC003937 (PI: Svirsky), and by a research contract from Cochlear Ltd.

